# Using Deep Learning Models of Gene Regulation to Guide Drug Prioritization

**DOI:** 10.64898/2026.05.11.724354

**Authors:** Xiaoqin Huang, Ivan Ovcharenko

## Abstract

Drug repurposing offers a cost-effective strategy to accelerate therapeutic discovery, but most computational approaches fail to model noncoding genetic variation. Because over 90% of genome-wide association study (GWAS) risk variants reside in noncoding regions, linking regulatory variation to therapeutic hypotheses remains a major challenge. Here, we developed an integrative deep learning framework that links allele-specific enhancer prediction to transcription factor (TF)-centered gene expression changes and drug-induced transcriptional profiles to prioritize candidate therapeutics. Our cell type-specific deep learning enhancer models accurately distinguish active enhancers across seven cell lines. Using breast cancer as a proof-of-concept, we found that GWAS heritability is significantly enriched in MCF7 enhancers, supporting MCF7 as the cellular context for this disease. Allele-specific variant scoring identified breast cancer risk variants with strong allele-dependent effects, and attribution-based motif discovery revealed enrichment of FOXA1-associated motif features, consistent with FOXA1 upregulation in primary tumors. Integration of the FOXA1 knockdown-induced and drug-induced gene expression profiles identified 63 candidate compounds for treatment of breast cancer, including 18 approved drugs, with recovery of the known breast cancer therapy fulvestrant. Among prioritized compounds, 54% showed anti-correlated transcriptional effects across eight core breast cancer pathways, compared to 5.3% of non-prioritized compounds. Integration of drug-gene interaction data further refined these to eight compounds with supporting experimental or clinical evidence. Together, these results establish a regulatory variant-guided drug repurposing framework that connects noncoding genetic variation to therapeutic candidates and provides a generalizable strategy for translating the noncoding genome into pharmacologically relevant hypotheses.

## Background

The development of new therapeutics remains slow, costly, and characterized by high attrition rates [1–5]. Traditional drug discovery typically requires 10-17 years and costs over $2 billion per approved drug, with only approximately 11% of compounds entering clinical trials ultimately receiving approval, and success rates even lower in areas such as neurodegenerative diseases [6]. Drug repurposing offers a complementary strategy by identifying new indications for compounds with established safety profiles, thereby potentially accelerating therapeutic discovery while reducing cost and risk [7, 8]. A growing number of computational approaches have been developed to facilitate drug repurposing, including gene expression signature matching, network-based inference, and structure-based modeling [1, 9]. Many of these strategies leverage molecular phenotypes, such as drug-induced or disease-associated transcriptional profiles, to identify compounds with opposing or corrective effects, and have yielded promising results across diverse disease contexts, such as muscle atrophy [10], neurodegenerative disorders [11, 12], and infectious diseases [13]. However, most existing frameworks rely primarily on transcriptional anti-correlation or network similarity and lack explicit modeling of how genetic risk perturbs disease-relevant regulatory programs.

Genome-wide association studies (GWAS) have revealed that over 90% of risk variants for complex traits reside in noncoding regions of the genome, particularly within distal regulatory elements such as enhancers [14–16]. Enhancers orchestrate cell type-specific transcriptional programs through cooperative binding of TFs, and perturbation of enhancer activity represents a major mechanism by which genetic variation influences disease risk [17]. However, most GWAS-informed drug discovery approaches prioritize genes based on genomic proximity or statistical association without explicitly modeling allele-specific regulatory effects within enhancers or the TF-associated gene expression changes through which these variants influence disease risk [18, 19]. Consequently, how noncoding genetic risk variants can be translated into therapeutic hypotheses through their regulatory effects on gene expression remains poorly defined.

Recent advances in deep learning-based regulatory modeling enable prediction of enhancer activity directly from DNA sequence and inference of allele-specific effects of noncoding variants [20]. These models provide a framework for moving beyond static association signals toward identification of TF-centered gene expression changes associated with genetic risk [21–25]. We therefore hypothesize that disease-associated noncoding variants disproportionately affect enhancer sequence features associated with specific TFs, and that compounds whose drug-induced gene expression changes directionally concordant with TF-associated gene expression changes represent computationally supported candidates for therapeutic repurposing.

Here, we present an integrative framework that moves from noncoding genetic risk to therapeutic prioritization by modeling allele-specific enhancer prediction, inferring disease-relevant TFs, and matching TF knockdown-induced gene expression to drug-induced transcriptional responses. Our approach combines cell type-specific deep learning-based enhancer prediction, allele-specific variant impact assessment, and motif discovery to identify trait-relevant TFs, integration of TF knockdown-induced gene expression with drug-induced transcriptional responses to quantify directional concordance; and pathway-level anti-correlation and curated drug-gene interactions to refine candidate compounds. We illustrate this strategy using breast cancer as a proof-of-concept application, demonstrating how risk-associated enhancer sequence features preferentially converge on FOXA1-associated motifs and how compounds whose drug-induced gene expression changes directionally concordant with FOXA1 knockdown show anti-correlated transcriptional effects to core breast cancer-associated hallmark pathways. This proof-of-concept application establishes a transferable computational framework for regulatory variant-guided drug repurposing that can be extended to other complex traits as matched epigenomic, TF perturbation, and drug transcriptional datasets become available.

## Results

### Overview of the framework

The overall analytical framework is summarized in Fig. 1. We first defined cell type-specific enhancers by integrating chromatin accessibility (ATAC-seq or DNase-seq) and H3K27ac profiles across multiple cell lines obtained from ENCODE [26]. Cell type-specific enhancer models were trained using TREDNet, a two-phase deep learning framework [27]. Phase one was pre-trained on 4560 genomic and epigenomic profiles from ENCODE v4 [28], and phase two models were fine-tuned for each cell type using 2-kb enhancer and control sequences (see Methods). Using this model, GWAS variants were evaluated for allele-dependent differences in enhancer prediction, and candidate regulatory variants were identified based on allele-specific enhancer predictions. Attribution-based motif discovery (DeepLIFT [29] followed by TF-MoDISco [30]) was then applied to infer TF motif features preferentially associated with risk alleles.

**Figure 1.**
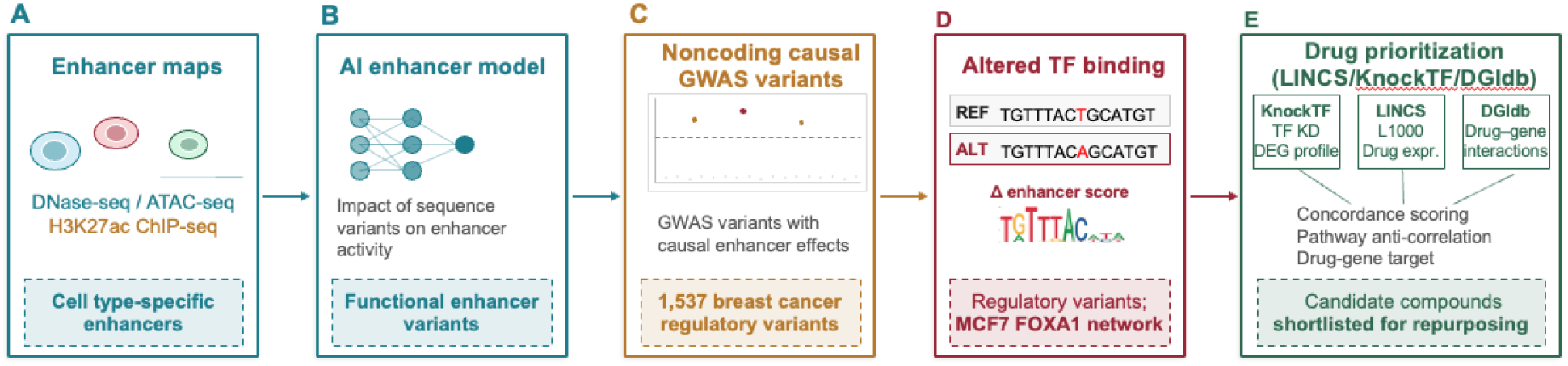
Regulatory variant-guided drug repurposing framework. (A) Cell type-specific enhancer definition from chromatin accessibility and H3K27ac profiles. (B) Cell type-specific enhancer models are trained using TREDNet. (C) Identification of candidate regulatory variants through allele-specific enhancer prediction. (D) Inference of TF-associated motifs using attribution-based motif discovery. (E) Integration of TF perturbation, drug-induced gene expression, and drug-gene interaction data for compound prioritization.

To connect these regulatory features to downstream gene expression, TF knockdown-induced transcriptional profiles are integrated with drug-induced gene expression data from the LINCS L1000 dataset [31]. Compounds are prioritized based on transcriptional concordance with TF perturbation profiles from KnockTF 2.0 [32], followed by pathway-level anti-correlation analysis and integration of curated drug-gene interaction data from DGIdb 5.0 [33] to refine candidate therapeutics.

### Deep learning enhancer modeling and heritability enrichment link cell lines to disease GWAS

Cell-specific enhancer models were trained using the TREDNet two-phase deep learning framework [27], achieving robust performance on chromosome-level held-out test sets (see Methods), with AUROC of 0.91 and AUPRC of 0.65 in MCF7 cell line (Fig. S1A-B). These results indicate reliable discrimination of enhancers from human open chromatin regions not overlapping enhancer sites, using a 1:10 positive-to-negative sampling ratio.

Significant heritability enrichment was observed for trait-cell line pairs consistent with known cell type identity. MCF7 enhancer annotations span 2.4% of HapMap3 SNPs yet account for 25.1% of breast cancer SNP heritability (10.5-fold enrichment; p = 4.1 × 10^−6^; one-sided z-test with block jackknife standard error; linkage disequilibrium score regression (LDSC) partitioned heritability regression [34]; Fig. S1C), indicating that breast cancer genetic risk is disproportionately concentrated within MCF7-specific regulatory elements, consistent with MCF7’s origin as a breast cancer-derived cell line [35]. Based on this enrichment and the availability of matched TF perturbation and drug transcriptional data, we selected the MCF7-breast cancer pair for subsequent allele-specific modeling, TF inference, and drug prioritization analyses, as a proof-of-principle model.

### Allele-specific TF convergence at breast cancer risk loci

Focusing on the MCF7 breast cancer context, 20,619 breast cancer-associated variants passing the genome-wide significance threshold (p < 5 × 10^−8^) from full GWAS summary statistics [36] were used. FUMA mapped these variants to 151 genomic risk loci. To summarize their broader genomic distribution, we also intersected them with a predefined GRCh38 European LD block map (https://github.com/jmacdon/LDblocks_GRCh38/blob/master/data/pyrho_EUR_LD_blocks.bed), which assigned 18,800 variants to 141 LD blocks, with many variants falling within the same block. Of the 20,619 significant variants, 830 variants overlapped MCF7 enhancers (1-kb windows). For each variant, we computed an allele-specific enhancer score difference (Δscore), defined as the difference in TREDNet-predicted enhancer probability between the risk and protective allele sequences centered on a 2-kb window (Δscore = P_*risk allele*_ − P_*protective allele*_; see Methods). Variants were retained as candidate regulatory variants if at least one allele exceeded the model-derived enhancer probability threshold (false positive rate < 10%) and |Δscore| ranked within the top 10th percentile of the empirical distribution across all evaluated variants. This filtering identified 1,537 variants (7.5%) exhibiting strong allele-dependent differences in predicted enhancer probability, with risk alleles associated with both increases and decreases in enhancer prediction score, consistent with the heterogeneous regulatory architecture of complex trait loci (Fig. S2C).To identify TF motifs preferentially binding by breast cancer risk alleles through allele-specific differences in enhancer prediction, we applied attribution-based motif discovery to the 1,537 candidate regulatory variants. DeepLIFT [29], a feature attribution method that estimates the contribution of each input base to a neural network prediction relative to a reference input, was used to compute nucleotide-resolution contribution scores from TREDNet predictions for each allele sequence. These contribution scores were then analyzed with TF-MoDISco [30], which identifies short high-contribution subsequences (“seqlets”) and clusters recurrent seqlets into TF motif classes. In this framework, DeepLIFT identifies nucleotide positions contributing to allelic differences in enhancer prediction, and TF-MoDISco summarizes these signals into interpretable TF-associated sequence patterns. Across the 1,537 candidate variants, this approach identified 32 TF motif classes with varying degrees of allelic enrichment in risk versus protective allele attribution profiles (Fig. S2A; Table S1). Broadly acting architectural factors, including CTCF and SP2, showed no significant allelic bias (enrichment fold=1.0 and 1.13, respectively), indicating comparable contributions to enhancer prediction in both allele contexts. In contrast, a subset of lineage-associated TFs showed preferential enrichment of motif signatures in risk allele attribution profiles. Most prominently, FOXA1-associated motif features were strongly enriched in risk alleles relative to protective alleles (40 vs. 0 seqlets in risk vs. protective alleles, p = 4.9×10^-12^), indicating that FOXA1 motif patterns contribute more strongly to TREDNet-predicted enhancer prediction in risk allele sequences. Related forkhead family members, including FOXD2, showed comparable directional enrichment (35 vs. 0 seqlets in risk vs. protective alleles). These results indicate that breast cancer risk alleles are disproportionately associated with forkhead family TF motifs.

We integrated model-derived allelic TF motif enrichment with tumor gene expression from TCGA-BRCA to further prioritize TFs. Of the 32 TF motif classes evaluated, 15 showed preferential enrichment in risk allele attribution profiles and 11 in protective allele profiles (Table S1). We first applied two data-driven criteria: (i) significant enrichment of TF motifs in risk versus protective allele attribution profiles, and (ii) concordant upregulation of the corresponding TF in primary breast tumors relative to matched normal tissue. Among TFs satisfying both criteria, FOXA1 and FOXD2 exhibited strong motif enrichment (40 vs. 0 and 35 vs. 0 seqlets in risk versus protective alleles, respectively) and high tumor-specific upregulation (log_2_ fold change (log_2_FC) = 2.18 and 2.26, respectively; p = 5.9×10^-167^ and 1.1×10^-69^, Table S1). In contrast, other TFs with highly enriched motifs, such as RXRG and BATF3, showed reduced expression in tumors (log_2_FC = −3.03 and −0.31, respectively; p = 9.8×10^-80^ and 1.9×10^-3^), indicating discordance between motif enrichment and transcriptional activity (Table S1). We next examined whether these prioritized TFs have reported roles in breast cancer regulatory programs. FOXA1 has been shown to bind enhancer regions and promote estrogen receptor recruitment in breast cancer cell models, thereby influencing estrogen receptor-dependent transcription [37–39]. Experimental modulation of FOXA1 in breast cancer models disrupts estrogen receptor-dependent transcription, alters enhancer accessibility, and reduces tumor cell proliferation [37, 38]. In contrast, little evidence currently links FOXD2 to breast cancer-related regulatory processes. To assess the disease relevance of FOXA1-regulated transcriptional programs, we evaluated overlap between genes differentially expressed upon FOXA1 knockdown in MCF7 cells and genes dysregulated in TCGA-BRCA tumors. FOXA1 perturbation affected the expression of 3,090 genes (false discovery rate (FDR) adjusted p < 0.05 from differential expression analysis, as reported in KnockTF 2.0, a curated database of TF knockdown and knockout gene expression profiles comprising 456 TFs across 357 human cell lines [32]), of which 205 (6.6%) overlapped with tumor-associated differentially expressed genes (odds ratio = 1.18; p = 0.024, one-sided Fisher’s exact test). Additionally, 202 FOXA1-regulated genes overlapped with genes mapped to breast cancer GWAS loci (FUMA [40]), supporting the relevance of FOXA1-regulated transcriptional programs to inherited disease risk. We therefore next focused drug prioritization on identifying compounds whose transcriptional effects are concordant with FOXA1 knockdown-induced gene expression changes.

### Drug candidate prioritization through FOXA1-centered transcriptional concordance

To identify such compounds, we integrated the FOXA1 knockdown-induced gene expression profile from the MCF7 cell line (3,090 differentially expressed genes; FDR adjusted p < 0.05 from differential expression analysis, as reported in KnockTF 2.0) with drug-induced transcriptional profiles from the LINCS L1000 dataset. For each compound in LINCS L1000, we computed a FOXA1 knockdown concordance score, defined as the mean drug-induced z-score of FOXA1 knockdown-upregulated genes minus the mean drug-induced z-score of FOXA1 knockdown-downregulated genes, such that positive values indicate transcriptional concordance with FOXA1 knockdown (see Methods). Candidate compounds were required to satisfy two criteria: (i) statistically significant enrichment between drug-induced gene expression changes and FOXA1 knockdown-associated differentially expressed gene sets, and (ii) a FOXA1 knockdown concordance score within the top 10th percentile across all evaluated compounds. Approved drug enrichment was maximal and statistically significant exclusively at the 90th percentile threshold (2.1-fold; odds ratio = 2.5; p = 0.002, one-sided Fisher’s exact test), declining and losing significance at more stringent or more lenient thresholds (Table S2). This filtering yielded 63 candidate compounds, of which 18 are approved for human clinical use by the FDA or equivalent regulatory agencies according to DrugCentral [41] (a curated database of international drug approval status) and ChEMBL [42] (a curated database of bioactive compounds with clinical development phase annotations), representing a 2.1-fold enrichment relative to the full LINCS L1000 library (odds ratio = 2.5; p = 0.002, one-sided Fisher’s exact test). To stratify candidates by clinical evidence, compounds were categorized into three tiers: Tier 1-approved drugs with registered breast cancer clinical trials in ClinicalTrials.gov [43]; Tier 2-approved drugs without breast cancer-specific trials; and Tier 3-investigational compounds (with or without breast cancer clinical trials).

Notably, our model has solidly recovered fulvestrant, an FDA approved drug for treating metastatic breast cancer in postmenopausal women [44]. To characterize transcriptional patterns among the 63 FOXA1-prioritized compounds, we computed enrichment of all available MCF7 TF knockdown gene sets from KnockTF 2.0 against drug-induced gene expression profiles. For each TF independently, we quantified directional concordance by measuring whether genes upregulated upon TF knockdown were also upregulated by the drug (up-up) and whether genes downregulated upon TF knockdown were also downregulated by the drug (down-down) (Fig. 2). This concordance framework was used because our goal was to identify compounds whose transcriptional effects mirror the expression pattern induced by TF knockdown, such that higher concordance indicates greater similarity between the drug-induced profile and the TF knockdown signature. Although compounds were selected based on FOXA1 concordance, multiple additional TF-associated gene sets showed consistent enrichment across prioritized compounds, indicating that these drugs may influence coordinated transcriptional programs beyond FOXA1 alone. Among TFs with significant directional concordant enrichment, TFAP2C-associated gene sets exhibited prominent up-up concordance (Fig. 2). TFAP2C has been shown to regulate gene expression associated with luminal breast epithelial cells and estrogen receptor signaling [45, 46], suggesting that prioritized compounds coordinately suppress both FOXA1- and TFAP2C-dependent transcriptional programs. In addition, gene sets associated with KDM3A and RARA showed significant down-down concordance across prioritized compounds. KDM3A encodes a histone H3K9 demethylase implicated in estrogen receptor-dependent transcription [47], while RARA encodes a nuclear receptor whose activity intersects with estrogen receptor signaling [48]. GATA3-associated gene sets showed significant down-down concordance across prioritized compounds, indicating that genes reduced after GATA3 knockdown were likewise reduced by drug treatment. Given that GATA3 is a central regulator of mammary luminal-cell differentiation and a hallmark factor in luminal breast cancer [49], this enrichment further supports that prioritized compounds influence coordinated luminal transcriptional programs beyond FOXA1 alone. The coordinated enrichment of multiple TF-associated gene sets indicates that the transcriptional effects of prioritized compounds are not limited to FOXA1-regulated transcriptional programs but extend to additional TF-linked gene expression signatures. Given that FOXA1 functions as a pioneer factor that establishes chromatin accessibility for estrogen receptor binding [39], ESR1-associated gene sets would be expected to show concordance among prioritized compounds; however, ESR1 concordance was weaker than anticipated (Fig. 2), which may reflect the requirement for ligand-bound receptor recruitment to FOXA1-primed chromatin sites-a cooperative dependency not captured in ligand-free TF knockdown datasets from KnockTF 2.0 [50].

**Figure 2.**
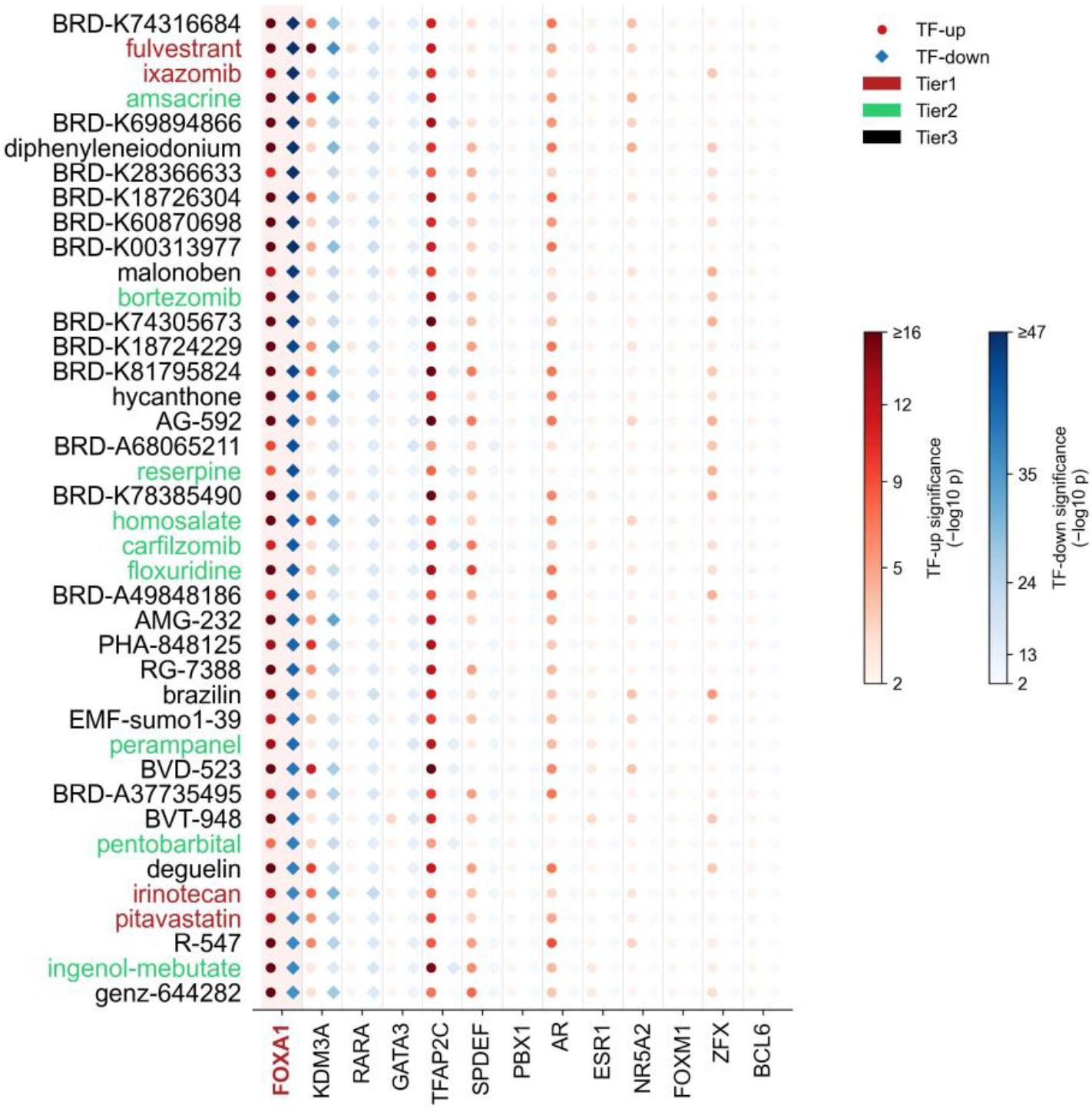
Directional transcriptional concordance between prioritized compounds and FOXA1 knockdown-induced gene expression. Heatmaps display the top 40 compounds ranked by FOXA1 knockdown concordance score whose drug-induced gene expression changes show directional concordance with FOXA1 knockdown-induced gene expression in MCF7. The x-axis shows all available MCF7 TF knockdown profiles (genes with FDR adjusted p < 0.05 from differential expression analysis, as reported in KnockTF 2.0); the y-axis shows compound names colored by candidate tier. Color intensity represents enrichment significance (-log_10_ p-value, one-sided Fisher’s exact test); with higher values indicating greater overlap.

Among the 63 prioritized compounds, the FDA-approved selective estrogen receptor degrader (SERD) fulvestrant ranked fourth among Tier 1 compounds by FOXA1 knockdown concordance score, behind ixazomib, pitavastatin, and pralatrexate (Fig. 3B). Fulvestrant degrades ESR1 protein at enhancers where FOXA1 establishes chromatin accessibility, thereby disrupting estrogen receptor-dependent transcription [37, 51]. Its recovery among the top Tier 1 compounds is therefore expected: the transcriptional consequences of ESR1 degradation-suppression of estrogen receptor-dependent gene expression, are directionally concordant with FOXA1 knockdown-induced gene expression changes, as both converge on the same breast cancer transcriptional axis. The recovery of fulvestrant within the top Tier 1 compounds supports the biological validity of FOXA1-centered transcriptional concordance as a prioritization criterion. Among compounds with higher concordance scores, ixazomib and bortezomib-both proteasome inhibitors, ranked at the top of the candidate list (Fig. 3B), highlighting the capacity of the framework to identify pharmacologically diverse mechanisms beyond the estrogen receptor axis.

**Figure 3.**
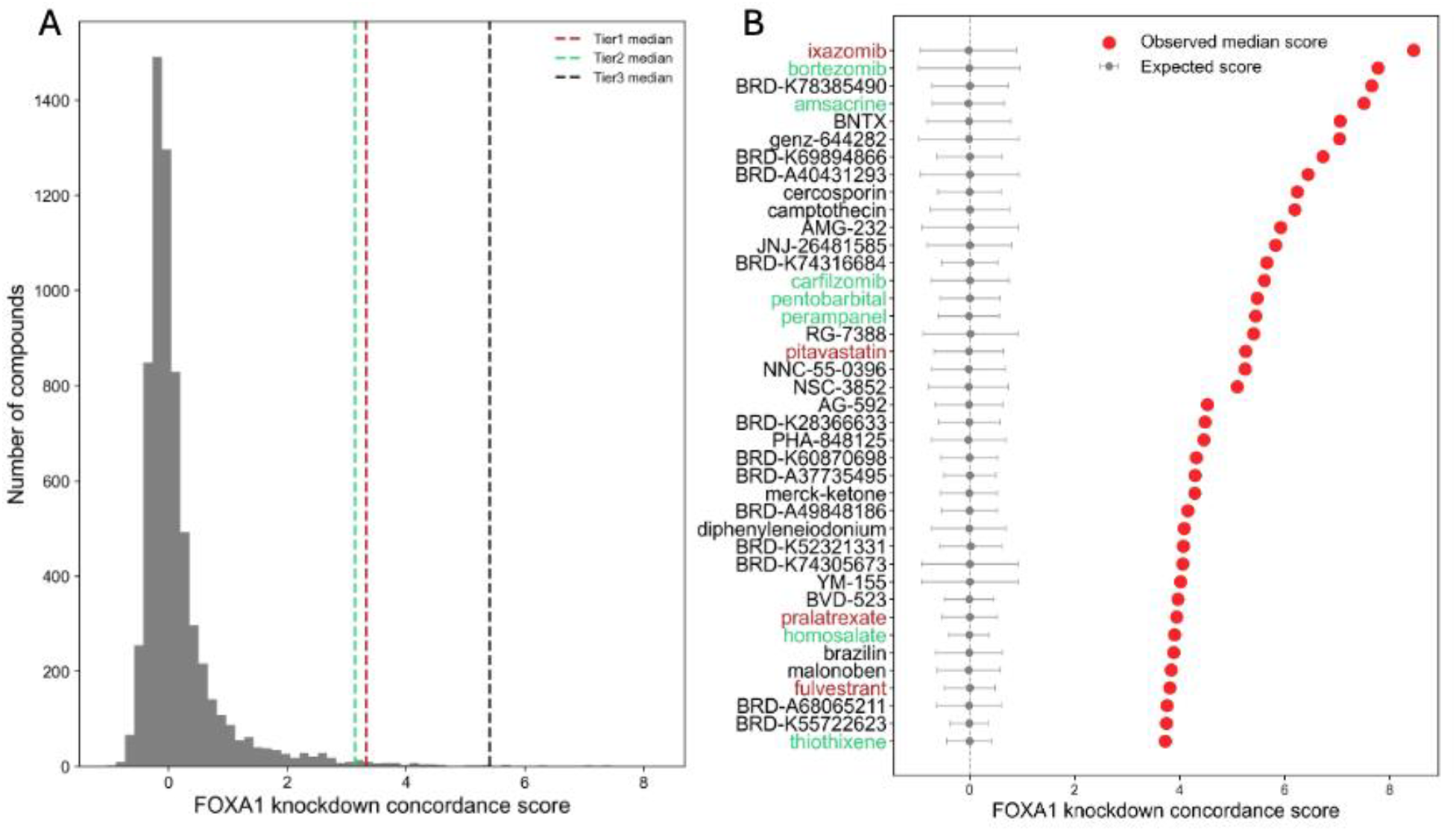
FOXA1 knockdown concordance scores quantify directional concordance between drug-induced and FOXA1 knockdown-induced gene expression. (A) Distribution of FOXA1 knockdown concordance scores across all 6,605 compounds. Dotted vertical lines indicate median concordance scores for Tier 1 (red), Tier 2 (green), and Tier 3 (black) compounds. (B) Observed and expected FOXA1 knockdown concordance scores for the top 40 prioritized compounds. Compounds are ranked by the observed median FOXA1 knockdown concordance score across LINCS signatures in MCF7 cells (red points). Gray points and horizontal error bars indicate the permutation-derived expected score (null mean ± SD), estimated from random gene sets matched to the sizes of the FOXA1 knockdown-upregulated and knockdown-downregulated gene sets. Colored compound names indicate tier categories.

To assess directional concordance between drug-induced transcriptional responses and FOXA1 suppression, we computed a FOXA1 knockdown concordance score for each compound across the full LINCS L1000 library treated in MCF7 (N = 6,605 compounds; median = −0.02; IQR = 0.48; Fig. 3A). The 63 prioritized compounds exhibited substantially higher concordance scores than the full LINCS background (median = 4.01; IQR = 1.84), with all three tiers significantly exceeding the background distribution (Tier 1: median = 3.94, p = 6.99 × 10^−5^; Tier 2: median = 3.73, p = 4.77 × 10^−10^; Tier 3: median = 4.06, p = 2.98 × 10^−30^; one-sided Mann-Whitney U test; Fig. 3A). Across all 63 candidates combined, concordance scores were significantly elevated relative to the full compound library (p = 1.96 × 10^−41^; one-sided Mann-Whitney U test), suggesting that the selection criteria effectively enriched for compounds with transcriptional concordance with FOXA1 suppression. Concordance scores did not differ significantly across tiers (Tier 1 vs Tier 2: p = 0.25; Tier 1 vs Tier 3: p = 0.35; Tier 2 vs Tier 3: p = 0.69; Mann-Whitney U test), consistent with the tier classification reflecting clinical evidence rather than transcriptional concordance strength. Permutation testing indicated that observed knockdown concordance scores of all 63 compounds significantly exceeded expected values which were the mean values of the empirical null distribution derived from 5,000 random gene set permutations (p < 2.0 × 10^−4^ for all compounds; permutation null mean ranging from −0.024 to 0.021; one-sided permutation test; Fig. 3B), demonstrating that directional concordance between drug-induced and FOXA1 knockdown-induced gene expression changes is statistically robust and consistent across prioritized candidates.

### Prioritized compounds exhibit transcriptional signatures anti-correlated with breast cancer-associated pathways

Breast tumors are characterized by transcriptional programs associated with proliferation, cell-cycle progression, oncogenic growth signaling and hormone-responsive signaling [52]. In breast cancer, pathways such as E2F targets [53], G2M checkpoint [54], MYC targets [55], mTORC1 signaling [56], unfolded protein response [57], and estrogen response [58] have been associated with aggressive clinical features, metastasis risk, and endocrine-responsive biology. To evaluate whether the transcriptional effects of prioritized compounds are anti-correlated with disease-associated pathway activity, we defined breast cancer-associated transcriptional programs using differential gene expression analysis of primary tumors versus matched normal tissue, followed by hallmark pathway enrichment analysis (Fig. 4A). These pathways were likewise among the canonical programs enriched in tumors.

**Figure 4.**
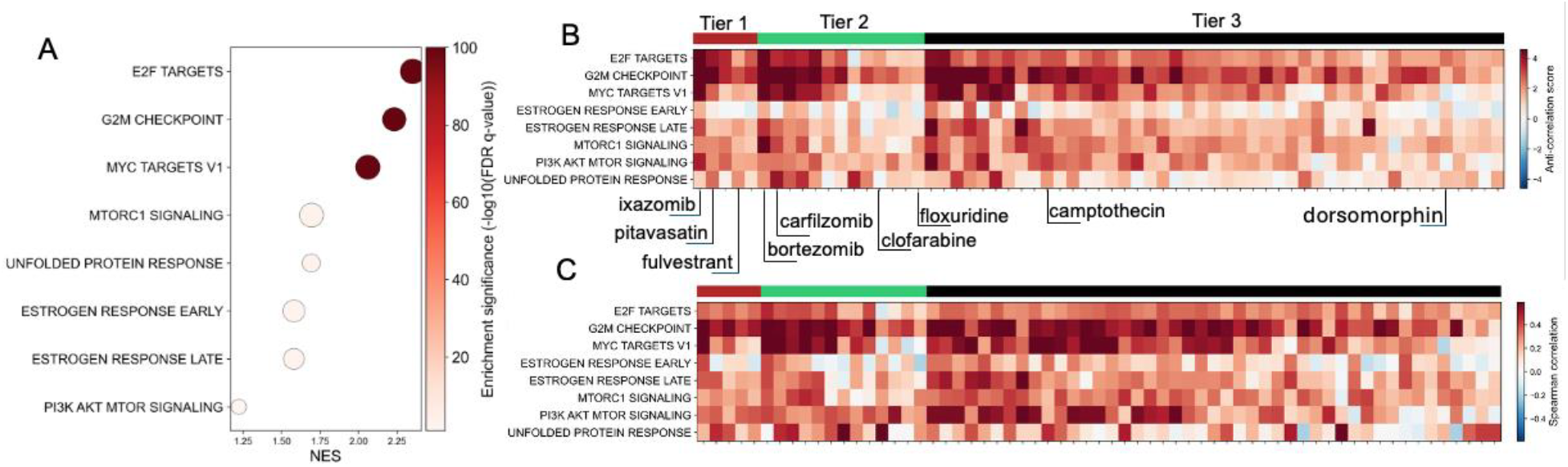
Prioritized compounds show transcriptional effects anti-correlated to core breast cancer-associated hallmark pathway activity. (A) Hallmark pathway enrichment in TCGA-BRCA differentially expressed genes (primary tumors versus matched normal breast tissue; DESeq2, FDR adjusted p < 0.01, |log_2_FC| > 2). NES, normalized enrichment score; positive values indicate enrichment among tumor-upregulated genes. (B) Pathway-level anti-correlation scores for 63 prioritized compounds across eight breast cancer-associated hallmark pathways. Color scale is centered at zero and capped at the 95th percentile of absolute values for visualization. (C) Rank-based directional concordance measured by Spearman correlation (ρ) between drug-induced gene expression changes and a signed disease direction vector (+1 for tumor-upregulated genes and −1 for tumor-downregulated genes within each pathway). Positive values indicate transcriptional changes anti-correlated to tumor-associated gene expression. Compound labels are colored by candidate tier. For readability, only representative compounds from each tier are labeled; the full compound list is provided in Supplementary Table S3.

To quantify the extent to which prioritized compounds are transcriptionally anti-correlated with breast cancer-associated pathway activity, we computed two complementary metrics for each of the 6,605 LINCS compounds: an anti-correlation score, defined as the mean drug-induced z-score of tumor-downregulated pathway genes minus the mean z-score of tumor-upregulated pathway genes (positive values correspond to anti-correlated transcriptional changes to the tumor expression pattern), and a rank-based Spearman correlation between drug-induced z-scores and a signed disease direction vector (positive values indicating directional anti-correlation; see Methods; Fig. 4B-C). Among the 63 FOXA1-prioritized compounds, 34 (54%) exhibited positive anti-correlation scores across all eight evaluated hallmark pathways simultaneously, compared to 5.3% of the 6,542 non-prioritized LINCS compounds (10.3-fold enrichment; p = 2.3 × 10^−26^, one-sided Fisher’s exact test). Consistent results were obtained using the Spearman metric (54% vs 4.1% of non-prioritized compounds; 13.1-fold enrichment; p = 1.3 × 10^−29^), indicating that the enrichment is consistent across scoring approaches. An additional 23 compounds (37%) showed positive scores in seven of eight pathways, and the remaining six compounds (9%) in six of eight pathways, indicating that the majority of prioritized compounds exhibit transcriptional changes anti-correlated to core breast cancer pathway activity.

Cell proliferation-associated pathways showed the most consistent anti-correlated transcriptional changes across compounds: all 63 compounds exhibited positive scores for the G2M checkpoint (100%), 62 for E2F targets (98.4%), and 59 for MYC targets (93.6%; Fig. 4B). Pathways related to growth and stress responses also showed similar patterns-mTORC1 signaling in 62 compounds (98.4%), PI3K/AKT/mTOR signaling in 60 (95.2%), and the unfolded protein response in 57 (90.5%). For hormone-responsive programs, 62 compounds (98.4%) showed positive scores for the estrogen response late signature, whereas a smaller subset (44 of 63; 69.8%) showed positive scores for the estrogen response early signature. This difference indicates that anti-correlated transcriptional changes are more consistently observed for late estrogen response genes than for early response genes [37]. Comparable results were obtained using the Spearman metric across all pathways (Fig. 4C, Table S4), indicating consistent patterns across scoring approaches.

Collectively, 57 of 63 prioritized compounds (91%) showed positive scores in at least seven of eight core breast cancer hallmark pathways, indicating that FOXA1-centered prioritization is associated with compounds whose transcriptional effects anti-correlated with multiple disease-associated pathway signatures, supporting their prioritization for further evaluation.

### Curated drug-gene interactions provide independent support for prioritized candidates

To determine whether the known molecular targets of prioritized compounds overlap with genes differentially expressed upon FOXA1 knockdown in MCF7, we cross-referenced each compound against DGIdb 5.0 [33], a publicly accessible database that aggregates drug-gene interaction data from multiple curated sources including ChEMBL [42], PharmGKB [59], and CIViC [60], and catalogs inhibitory, activating, and other experimentally supported interaction types between drugs and their molecular targets [33]. We then assessed whether documented drug targets overlapped with the 3,090 genes differentially expressed upon FOXA1 knockdown in MCF7 (FDR adjusted p < 0.05 from differential expression analysis, as reported in KnockTF 2.0). Among the 63 prioritized compounds, interactions between 13 compounds and genes differentially expressed upon FOXA1 knockdown were previously documented in DGIdb 5.0 [33], spanning Tier 1, Tier 2, and Tier 3 candidates (Fig. 5). These interactions provide independent support-derived from curated drug-target databases rather than transcriptional concordance scoring for the association between prioritized compounds and FOXA1-regulated gene expression.

**Figure 5.**
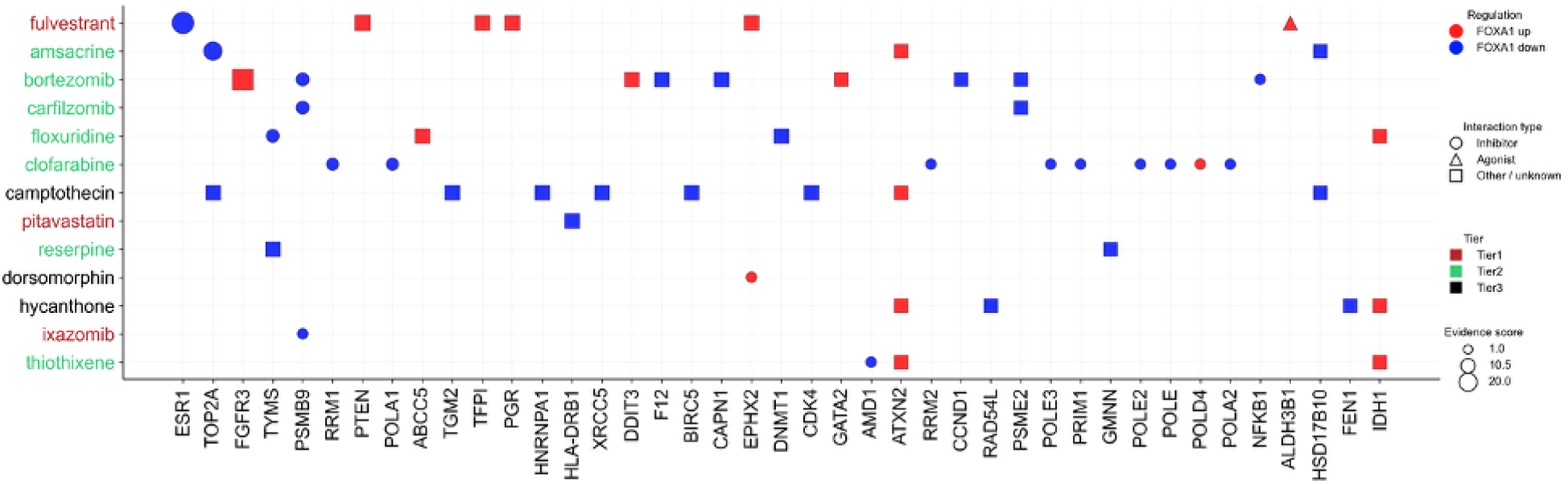
Drug-gene interactions for prioritized compounds and FOXA1 knockdown-associated genes. Bubble plot showing documented drug-gene interactions from DGIdb 5.0 for 13 of 63 prioritized compounds with at least one interaction partner among genes differentially expressed upon FOXA1 knockdown in MCF7 cells (KnockTF 2.0; FDR adjusted p < 0.05). Each marker represents an interaction between a compound (y-axis) and a FOXA1 knockdown-regulated gene (x-axis). Marker size corresponds to the DGIdb evidence score supporting the interaction, marker shape indicates the annotated interaction type, and marker color indicates direction of FOXA1 regulation (upregulated or downregulated upon knockdown). Compound names are colored by candidate tier.

Among Tier 1 compounds, fulvestrant has documented interactions with ESR1, PGR, PTEN, and ALDH3B1 in DGIdb 5.0 [33]. ESR1, for which fulvestrant has a documented inhibitory interaction, is among the genes downregulated upon FOXA1 knockdown in MCF7, indicating that fulvestrant’s known target is represented within the FOXA1-regulated gene set. Whether this overlap reflects direct ESR1 inhibition by fulvestrant, indirect effects mediated through FOXA1-dependent regulatory activity, or a combination of both cannot be determined from the present computational analysis.

Among Tier 2 compounds, the proteasome inhibitors bortezomib, carfilzomib, and ixazomib have documented interactions with PSMB9 in DGIdb 5.0 [33]; PSMB9 is among the genes downregulated upon FOXA1 knockdown in MCF7, indicating that these compounds share a documented target gene with FOXA1-regulated gene expression, though the functional significance of this overlap requires experimental validation. Pitavastatin has a documented interaction with HLA-DRB1 [61, 62], which is also among genes differentially expressed upon FOXA1 knockdown in MCF7, consistent with a potential link between its immunomodulatory effects and FOXA1-regulated transcriptional changes. Clofarabine and floxuridine have documented interactions with multiple genes differentially expressed upon FOXA1 knockdown. For example, clofarabine interacts with RRM1, RRM2, and POLA1-DNA replication-associated genes downregulated upon FOXA1 knockdown-indicating overlap between its known targets and FOXA1-regulated gene expression. Although some individual drug-gene interactions exhibit directionality discordant with the FOXA1 knockdown-associated expression changes, the mechanistic basis of such discordance cannot be determined from curated interaction data alone and may reflect cell type-specific, condition-specific, or database annotation differences [33].

Among Tier 3 compounds, camptothecin and dorsomorphin have documented interactions with genes differentially expressed upon FOXA1 knockdown in MCF7; however, the interaction type for these compounds is not specified in the DGIdb 5.0, limiting interpretation of interaction directionality. Drug-gene interaction data were not used as a quantitative selection criterion but were incorporated as an independent qualitative filter, derived from curated drug-target databases rather than transcriptional data, alongside transcriptional concordance scoring and pathway-level anti-correlation analysis in the final compound shortlisting.

Combining transcriptional concordance scoring, pathway-level anti-correlation analysis, and curated drug-gene interaction data with clinical evidence, we derived a shortlist of eight candidate compounds (Table 1): ixazomib, pitavastatin, bortezomib, carfilzomib, floxuridine, clofarabine, camptothecin, and dorsomorphin. These compounds collectively show high FOXA1 knockdown concordance scores, transcriptional effects that are anti-correlated with core breast cancer-associated hallmark pathways across scoring metrics, and documented interactions with FOXA1-regulated genes. As a proof-of-concept application in breast cancer, these results demonstrate that integrating allele-specific enhancer modeling, TF-centered gene expression analysis, and curated drug-target data can prioritize candidate compounds linked to noncoding genetic risk. This framework can be extended to other complex traits and cellular contexts as matched epigenomic, perturbation, and drug transcriptional datasets become available.

**Table 1.** Final prioritized drug candidates with transcriptional concordance and literature support. Binary values (1/0) indicate presence or absence of approval status (any indication) and breast cancer clinical trial annotation.

| Drug | FOXA1<br>KD<br>concordance<br>score | Fraction of<br>positive<br>pathway<br>opposition | Approved<br>(any<br>indication) | Breast<br>cancer<br>clinical<br>trial | Supporting literature evidence |
| --- | --- | --- | --- | --- | --- |
| Ixazomib | 8.45 | 1 | 1 | 1 | Ixazomib in combination with fulvestrant demonstrated a favorable safety profile and antitumor activity in patients with fulvestrant-resistant, advanced estrogen receptor-positive breast cancer. (NCT02993094; https://doi.org/10.1002/onco.13733) |
| Pitavastatin | 5.26 | 1 | 1 | 1 | Pitavastatin was reported to overcome paclitaxel resistance in triple-negative breast cancer models. (NCT04705909; PMID: 41126317) |
| Bortezomib | 7.78 | 1 | 1 | 0 | Bortezomib has been shown to inhibit breast cancer growth in preclinical studies. (PMID: 20843837) |
| Carfilzomib | 5.61 | 0.875 | 1 | 0 | An albumin-coated nanocarrier formulation of carfilzomib demonstrated improved metabolic stability and enhanced cytotoxic effects in breast cancer cells in vitro. (PMID: 30954620) |
| Floxuridine | 3.24 | 0.875 | 1 | 0 | Camptothecin-floxuridine conjugate nanocapsules enhanced anti-metastatic efficacy in breast cancer models. (https://pubs.acs.org/doi/10.1021/acsami.8b11723) |
| Clofarabine | 3.14 | 0.875 | 1 | 0 | Clofarabine exhibits potent anti-breast cancer activity, although cytotoxicity toward normal cells has been reported; modified analogs have shown improved selectivity. (https://doi.org/10.1016/j.bmcl.2025.130349) |
| Camptothecin | 6.20 | 1 | 0 | 0 | Tumor-targeted nanocrystal formulations of camptothecin demonstrated improved therapeutic efficacy in breast cancer models. (https://doi.org/10.1016/j.xphs.2025.103951) |
| Dorsomorphin | 3.14 | 0.75 | 0 | 0 | Dorsomorphin has been reported as an inhibitor of dickkopf-1 (DKK1), with implications for breast cancer progression. (PMID: 32001001) |

## Discussion

In this study, we developed an integrative computational framework that connects disease-associated noncoding variants to drug repurposing candidates by linking allele-specific enhancer prediction differences to disease-associated transcriptional changes and drug-induced transcriptional responses, using FOXA1-centered gene expression changes in breast cancer as a proof-of-concept application. We show that candidate regulatory variants preferentially alter enhancer prediction scores at loci enriched for FOXA1-associated motif features. In addition, compounds whose transcriptional effects are concordant with FOXA1 knockdown-induced gene expression exhibit anti-correlated transcriptional changes to core breast cancer-associated hallmark pathways. Together, these findings establish a computational link between noncoding genetic risk, TF-centered gene expression, and pharmacologically relevant candidate compounds.

A central biological finding from our analysis is the convergence of breast cancer risk variants on FOXA1-associated motif features. FOXA1 functions as a pioneer TF that establishes accessible chromatin states in breast cancer and coordinates estrogen receptor-dependent transcriptional programs [39]. Aberrant FOXA1 activity has been implicated in endocrine-resistant and metastatic ER-positive breast cancer [38, 63]. In our analysis, risk alleles showed strong preferential enrichment of FOXA1-associated motif features among 32 evaluated TF motifs (40 vs. 0 seqlets in risk vs. protective alleles), and FOXA1 showed strong upregulation among risk allele-enriched TFs in TCGA-BRCA primary tumors (log_2_FC = 2.18; q-value = 5.9 × 10^−167^, DESeq2 Wald test). Together, these observations support a model in which dispersed noncoding variants converge on lineage-defining enhancer-associated regulatory programs linked to breast cancer-relevant transcriptional changes.

Importantly, TF motifs identified through TF-MoDISco reflect motif features prioritized by the enhancer prediction model based on attribution scores, rather than direct evidence of differential TF binding in vivo between alleles [29]. Enrichment of a motif in one allele therefore indicates a stronger model-predicted regulatory contribution within that sequence context but does not imply absence of binding capacity or functional engagement in the alternate allele. This distinction is particularly relevant for pioneer factors such as FOXA1, which often bind broadly across accessible chromatin landscapes. Accordingly, our interpretation emphasizes allele-dependent differences in model-predicted regulatory contribution rather than strict presence-absence of binding events.

While FOXA1 emerged as the strongest convergent factor based on allele-specific motif enrichment, gene expression in primary breast tumors, and functional evidence, enhancer activity is typically governed by cooperative binding of multiple TFs, with regulatory output arising from their combined interactions rather than a single dominant factor [64]. In addition to FOXA1-associated motif enrichment, downstream drug perturbation analyses revealed coordinated changes in gene expression associated with additional TF-regulated gene sets, including TFAP2C-, RARA-, and KDM3A-associated gene expression changes. TFAP2C, RARA, and KDM3A have established roles in estrogen receptor-dependent and breast cancer-associated gene expression [46, 47, 65], suggesting that prioritized compounds modulate a broader set of co-regulatory TF-associated transcriptional programs beyond FOXA1 alone. Within such a combinatorial enhancer framework, discordance between allele-specific motif enrichment and gene expression in primary breast tumors for certain TFs, such as RXRG and BATF3, is biologically plausible. For example, RXRG functions as a tumor suppressor frequently silenced in breast cancer, whereas BATF3 is primarily associated with immune cell regulation [48, 66, 67]. Motif enrichment may therefore reflect variant-driven changes in local sequence context that alter the relative contribution of individual TFs to enhancer prediction, without necessarily changing their bulk expression levels in tumor tissue. This pattern is consistent with the principle that multiple TFs cooperatively regulate enhancer activity, such that changes in individual TF contributions do not require concordant changes in bulk expression levels [68, 69]. Accordingly, drug prioritization was guided by directional concordance with FOXA1 knockdown-induced gene expression changes, reflecting modulation of a central regulatory axis rather than inhibition of FOXA1 in isolation.

Rather than directly identifying compounds that bind or inhibit a single TF, our framework prioritizes drugs whose transcriptional effects are directionally concordant with TF knockdown-induced gene expression changes in the corresponding cell line (e.g. FOXA1 in MCF7 cells). The recovery of fulvestrant within the top four Tier 1 compounds is consistent with the established co-dependency of estrogen receptor transcriptional activity on FOXA1-mediated chromatin accessibility [37], although whether this reflects direct ESR1 inhibition or indirect effects through FOXA1-dependent regulation requires experimental investigation. Beyond established therapies, several prioritized compounds have independent experimental or clinical evidence supporting their relevance in breast cancer. Ixazomib has shown antitumor activity in fulvestrant-resistant, advanced breast cancer in clinical studies [70], consistent with its high FOXA1 knockdown concordance score and transcriptional effects anti-correlated to all eight evaluated hallmark pathways in our analysis. Pitavastatin has been reported to overcome chemoresistance in breast cancer models [71], supporting its identification as a biologically supported repurposing candidate. Proteasome inhibitors such as bortezomib and carfilzomib have shown preclinical antitumor activity in breast cancer systems [72, 73]. Together, these examples indicate that FOXA1-centered transcriptional concordance scoring identifies compounds with prior clinical or experimental evidence in breast cancer. It also highlights additional candidates, such as pitavastatin and the antimetabolites floxuridine and clofarabine, warranting further experimental evaluation.

Methodologically, this work extends regulatory deep learning approaches beyond variant annotation toward drug repurposing. Previous drug repurposing frequently relied on global transcriptional anti-correlation or pathway overlap without explicit modeling of upstream regulatory perturbation. By integrating enhancer-based allele-specific modeling, TF knockdown-induced gene expression profiles, directional transcriptional concordance scoring, and pathway-level anti-correlation analysis, our framework incorporates upstream regulatory context into compound prioritization. This stepwise integration provides a quantitative strategy for translating noncoding genetic architecture into computationally supported therapeutic hypotheses.

This framework can be generalized to any disease with related matched data available. We have explored several other cell lines, and we obtained decent performance for cell specific enhancer models and identified paired disease - cell line. Of 50 cell lines with overlapping LINCS and KnockTF coverage, seven retained sufficient epigenomic annotation after excluding cell lines with incomplete chromatin accessibility or H3K27ac profiles, or with documented prior perturbations including chemical treatment or physical manipulation such as ultrasonic treatment: A673, HCT116, HepG2, IMR90, K562, MCF7, and PC3 (see Methods). For each cell line, we defined enhancers by intersecting chromatin accessibility peaks with H3K27ac signals, yielding between 29,000 and 98,000 cell type-specific enhancers per cell line. These enhancers served two purposes: training cell type-specific enhancer prediction models and constructing annotations for heritability enrichment analysis. Enhancer prediction models were trained using the TREDNet two-phase deep learning framework [27], achieving robust performance on chromosome-level held-out test sets (see Methods), with AUROC of 0.91-0.98 and AUPRC of 0.58-0.86 across all seven cell lines (Fig. S1A-B). These results indicate reliable discrimination of enhancers from human open chromatin regions not overlapping enhancer sites, using a 1:10 positive-to-negative sampling ratio. These enhancers were then formatted as 1-kb window annotations for LDSC partitioned heritability analysis, which regresses per-SNP chi-square statistics on annotation-specific LD scores to estimate whether variants within a given annotation explain a greater proportion of trait heritability than expected from genome-wide background LD structure [34]. We applied LDSC to 135 GWAS summary statistics representing 42 traits selected based on their relevance to the seven cell lines analyzed. For each trait, all available GWAS full summary statistics in the GWAS catalog were included, resulting in multiple datasets per trait. Significant heritability enrichment was observed for trait-cell line pairs consistent with known cell type identity. Following the rest steps of this framework, potential therapeutic compounds can be prioritized for those diseases as well.

Several limitations warrant consideration. First, inference of candidate regulatory variants and compound prioritization are computational and require experimental validation to confirm regulatory and therapeutic effects. Second, applicability of the framework is constrained by the availability and overlap of public epigenomic, TF perturbation, and drug-induced transcriptional datasets, limiting analyzable cellular contexts. Third, drug-gene interaction databases (e.g. DGIdb 5.0 [33]) are incomplete and not cell type-specific, which may obscure context-dependent regulatory relationships. Fourth, the use of cancer-derived cell lines introduces somatic alterations that may not fully reflect germline regulatory mechanisms. Additionally, enhancer prediction models rely on available epigenomic annotations and may not fully capture regulatory dynamics under disease-specific conditions. These models predict enhancer probability rather than direct enhancer activity, and allele-specific differences in prediction scores do not constitute direct evidence of differential regulatory function *in vivo*.

Future work should prioritize experimental validation of prioritized regulatory variants and compounds, expansion of TF perturbation and drug transcriptional datasets across diverse cellular systems, and application of enhancer-informed regulatory modeling to non-cancer complex traits. Extending this framework to genetically driven neurological, immunological, and metabolic disorders may enable systematic identification of context-specific regulatory vulnerabilities. More broadly, these findings support the potential of enhancer-centered regulatory modeling as a foundation for translating noncoding genetic risk into computationally informed therapeutic hypotheses, pending validation across additional traits and cellular contexts.

## Conclusions

In this study, we developed a regulatory variant-guided drug repurposing framework that connects noncoding genetic variation to candidate therapeutics by linking allele-specific enhancer modeling, TF-centered gene expression, and drug-induced transcriptional responses. Using breast cancer as a proof-of-concept, we show that risk-associated enhancer sequence features converge on FOXA1-associated regulatory programs and that compounds selected based on transcriptional concordance with FOXA1 perturbation exhibit anti-correlated signatures with disease-associated pathway activity. Integration of curated drug-gene interaction data further refines these candidates and highlights compounds with supporting experimental or clinical evidence. Together, these results demonstrate that modeling enhancer-level regulatory effects provides a systematic approach for linking noncoding genetic risk to candidate therapeutics. As integrative functional genomics resources continue to expand, this approach offers a scalable path for applying regulatory modelling to other complex traits.

## Methods

### Data collection

#### Drug-induced gene expression profiles

Phase I and II L1000 gene expression profiles (978 landmark genes) were obtained from the Broad Institute LINCS project [31] via GEO (GSE70138 and GSE92742). Drug-induced gene expression z-score matrices were used for downstream analyses.

#### TF knockdown-induced gene expression profiles

TF knockdown-induced gene expression profiles were retrieved from the KnockTF 2.0 database [32], comprising perturbations of 456 TFs across 357 human cell lines. Up- and down-regulated gene sets (adjusted p < 0.05 from differential expression analysis, as reported in the database KnockTF 2.0) were extracted for each TF-cell line pair.

#### Epigenomic profiles

Chromatin accessibility (DNase-seq or ATAC-seq) and enhancer-associated histone modification profiles (H3K27ac) were obtained from ENCODE [26]. Fifty cell lines overlapping between LINCS and KnockTF datasets were initially considered. After excluding cell lines with incomplete chromatin accessibility or H3K27ac profiles, or with documented prior perturbations including chemical treatment or physical manipulation such as ultrasonic treatment, seven cell lines were retained: A673, HCT116, HEPG2, IMR90, K562, MCF7 and PC3.

#### GWAS summary statistics

GWAS summary statistics for 135 studies spanning 42 traits were downloaded from the GWAS Catalog. Traits were selected based on known biological association with the cell line of origin (e.g., breast cancer traits paired with MCF7, blood traits paired with K562). All summary statistics were harmonized to the GRCh38 reference genome.

### Cell type-specific enhancer modeling

Enhancer prediction was performed using the TREDNet two-phase deep learning framework [27]. Phase one was pretrained on 4,560 genomic and epigenomic profiles from ENCODE v4 [28]. Phase two models were fine-tuned separately for each cell type using 2-kb genomic regions centered on open chromatin peaks overlapping H3K27ac signals. Coding regions, promoter-proximal regions (<2 kb from transcription start sites), and ENCODE blacklisted regions were excluded [74].

Each model was trained using fivefold cross-validation with chromosome-level holdout to prevent information leakage. Negative control sequences were sampled at a 10:1 ratio from all human open chromatin regions not overlapping positive enhancer sites or blacklisted regions.

### GWAS enrichment in enhancer regions

Linkage disequilibrium score regression (LDSC) [75] was used to evaluate enrichment of GWAS heritability within cell type-specific enhancer regions. Enhancer annotations were defined using 1-kb windows centered on predicted enhancer peak summits and annotated using the 1000 Genomes Project reference panel (hg38). GWAS summary statistics were processed in 500,000 SNP chunks using HapMap3 SNPs (w_hm3.snplist). Enrichment was assessed independently for each cell line. Statistical significance was determined from the regression coefficient estimated by LDSC using a z-test, with p < 0.05 considered significant.

### Allele-specific enhancer effect and motif inference

For breast cancer, which showed significant heritability enrichment within MCF7 enhancer annotations. For each variant, 2-kb sequences centered on the variant position were scored under both risk and protective alleles using the MCF7 enhancer model.

The allele-specific enhancer difference (Δscore) was defined as:

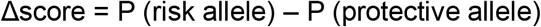

where P represents the predicted probability of the cell type-specific enhancer model for the allele-centered 2-kb sequence. Absolute Δscore values were used to quantify allele-specific differences in model-predicted enhancer probability. Candidate causal variants were defined using two criteria: (i) at least one allele exceeded the model-derived enhancer probability threshold corresponding to a false positive rate of 10%, and (ii) the absolute allele-specific prediction difference (|Δscore|) ranked within the top 10% of all evaluated GWAS variants (above the 90th percentile of the empirical |Δscore| distribution).

Attribution scores were computed using DeepLIFT [29, 76] with the negative enhancer sequence as background. TF-MoDISco [30] was applied to 500-bp windows centered on prioritized variants, importance-weighted subsequences, termed seqlets, representing recurrent sequence patterns within attribution score profiles (maximum 2,000 per metacluster).

### TF enrichment in drug perturbation profiles

For each TF-cell line pair with significant perturbation effects in KnockTF (adjusted p < 0.05 from differential expression analysis, as reported in the database KnockTF 2.0), up- and down-regulated gene sets were extracted. For each drug-cell line perturbation in LINCS, the top 250 up- and down-regulated genes were selected based on the z-score, consistent with standard LINCS L1000 analysis approaches [31].

Overlap between drug-induced gene sets and TF knockdown gene sets was assessed using Fisher’s exact test separately for up regulated genes upon TF knockdown in drug-induced upregulated genes (up-up) and down regulated genes upon TF knockdown in drug-induced downregulated genes (down-down) comparisons. TF enrichment p-values were computed for each drug-TF pair. For drugs with multiple perturbation profiles, the median enrichment p value was used.

### TF Knockdown concordance score

To quantify directional concordance between drug-induced and TF knockdown-associated transcriptional responses, a TF knockdown concordance score was calculated as:

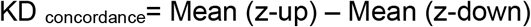

where Mean(z_up) and Mean(z_down) represent the average drug-induced z-scores for genes up-regulated and down-regulated in the TF knockdown-induced differentially expressed gene set, respectively. This formulation follows the connectivity score principle of the connectivity map framework [77], adapted to quantify directional concordance with TF knockdown-induced gene expression changes. For drugs with multiple perturbation profiles, the median concordance score across experiments was used. Statistical significance of the KD concordance score was assessed using an empirical permutation test that preserves the sizes of the TF knockdown up- and down-regulated gene sets. For each drug-TF pair, random gene sets of equal sizes (number of up- and down-regulated genes) were sampled without replacement from the L1000 landmark gene universe, and the concordance score was recomputed across 5,000 permutations to generate a null distribution. P-values were adjusted for multiple testing using the Benjamini-Hochberg FDR procedure.

Concordance score distributions were compared between prioritized compound tiers and the full LINCS background using one-sided Mann-Whitney U tests. Pairwise comparisons between tiers were performed using two-sided Mann-Whitney U tests.

### Drug prioritization

Candidate compounds were prioritized by requiring statistically significant directional enrichment in both up-up and down-down comparisons (Fisher’s exact test, p < 0.05) and a TF knockdown concordance score within the top 10th percentile across all evaluated LINCS compounds. The 10th percentile threshold was selected empirically based on maximization of approved drug enrichment across threshold values spanning the 75th to 95th percentile.

### Pathway-level anti-correlation analysis

For the breast cancer proof-of-concept analysis, differentially expressed genes (DEGs) were derived from TCGA-BRCA tumor versus matched normal RNA-seq data using TCGAbiolinks in R [78], with differential expression performed using DESeq2 (FDR adjusted p < 0.01, |log2 FC| > 2, Wald test). Hallmark pathway enrichment was assessed using preranked GSEA [79]. For each drug and pathway, we computed two complementary anti-correlation statistics: a raw score defined as the mean drug-induced z-score of disease-downregulated genes minus mean z-score of disease-upregulated genes, and a rank-based directional concordance score defined as the Spearman correlation between rank-transformed drug-induced z-scores and a signed disease direction vector. Positive values indicate transcriptional changes anti-correlated to the disease-associated expression pattern. This formulation follows the principle of gene expression anti-correlation described in the connectivity map framework [77] and uses a rank-based correlation approach analogous to methods employed in gene set enrichment analysis [79]. To contextualize these effects among prioritized compounds, the same metrics were computed for all 6,605 LINCS compounds treated in MCF7. Enrichment of compounds showing consistent anti-correlated transcriptional changes across all eight pathways was assessed by comparing the 63 prioritized compounds to the 6,542 non-prioritized compounds using a one-sided Fisher’s exact test. Statistical significance was further evaluated using an empirical permutation test in which drug-induced gene expression z-scores were randomly permuted across genes within each pathway gene set (n=5,000 permutations). P-values were adjusted for multiple testing using the Benjamini-Hochberg procedure.

### Drug annotation and drug-gene interaction analysis

Drug approval status and clinical trial annotations were obtained from DrugCentral [41] and ChEMBL [42] (accessed December 2025). Curated drug-gene interaction data were retrieved from DGIdb 5.0 [33] (accessed December 2025) and used as an independent source of drug-target evidence to assess overlap between documented drug targets and genes differentially expressed upon FOXA1 knockdown in MCF7.

## Supplementary Information

Supplementary materials are in the attachment.

## Declarations

## Acknowledgements

This research was supported by the Intramural Research Program of the National Institutes of Health (NIH), grant 1-ZIA-LM200881-12 (to I.O.). The contributions of the NIH author(s) are considered Works of the United States Government. The findings and conclusions presented in this paper are those of the author(s) and do not necessarily reflect the views of the NIH or the U.S. Department of Health and Human Services. In addition, this work utilized the computational resources of the NIH HPC Biowulf cluster (http://hpc.nih.gov).

## Authors’ contributions

X.H. conducted the experiments, performed the analyses, and wrote the manuscript. I.O. conceived and supervised the study, revised the manuscript, and provided critical feedback. All authors read and approved the final manuscript.

## Data availability

The pre-trained TREDNet model has been deposited at https://doi.org/10.5281/zenodo.8161621. Heritability partitioning analyses were performed using the standard S-LDSC implementation available at (https://github.com/bulik/ldsc). Human population variant data for heritability partitioning were sourced from the 1000 Genomes Project Phase 3, while baseline-LD model annotations (v2.2) were obtained from (https://doi.org/10.5281/zenodo.10515792). Breast cancer GWAS summary statistics were downloaded from the GWAS Catalog (accession GCST004988) at (https://www.ebi.ac.uk/gwas).

## Ethics approval and consent to participate

Not applicable.

