## supplementary figures and tables for "Using Deep Learning Models of Gene Regulation to Guide Drug Prioritization"

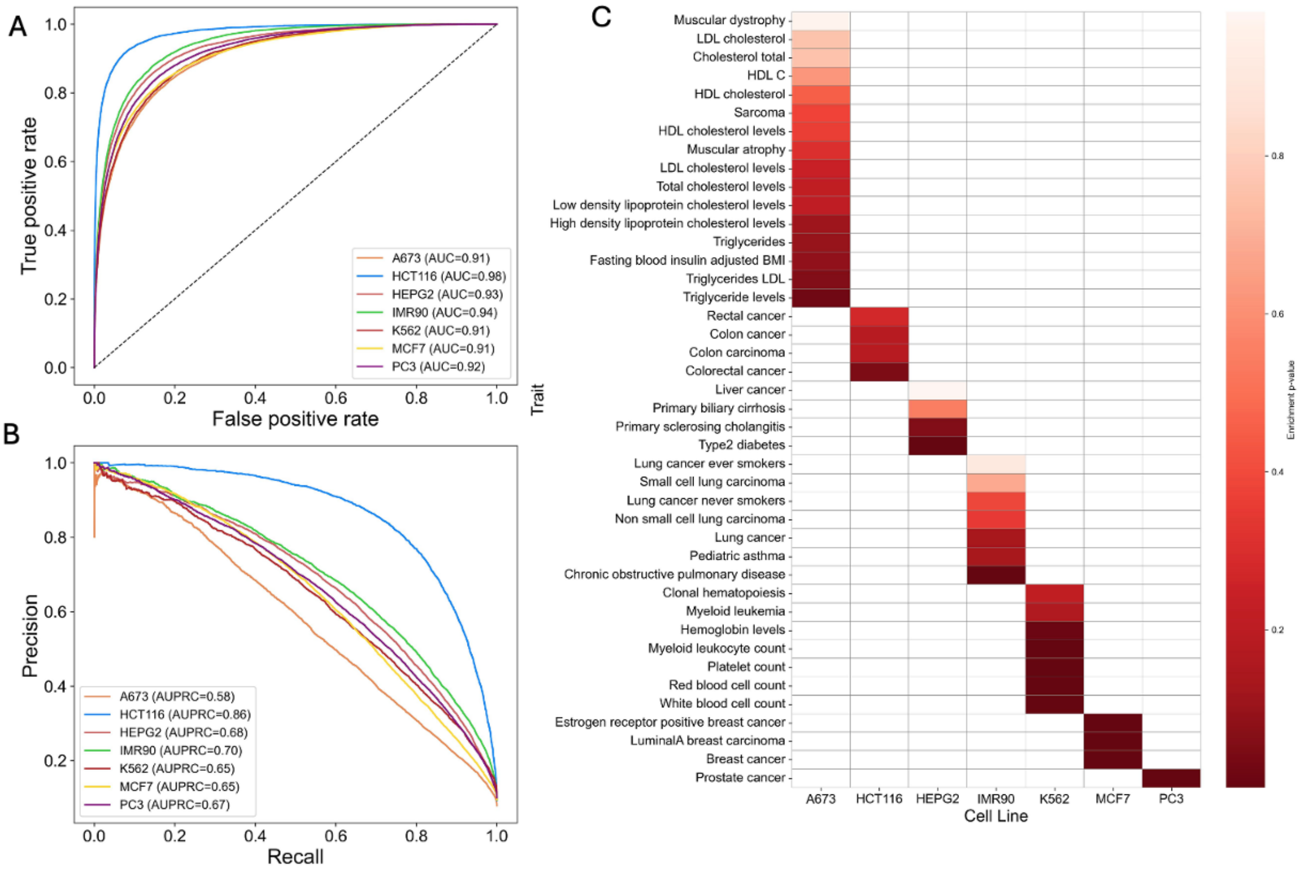


**Figure S1** Cell line-specific enhancer model performance and GWAS heritability enrichment. (A) Enhancer model performance (AUC) on the test set from a representative fold of chromosome-level cross-validation. (B) Enhancer model performance (AUPRC) on the test set from a representative fold of chromosome-level cross-validation. (C) GWAS heritability enrichment across cell line-specific enhancer annotations.

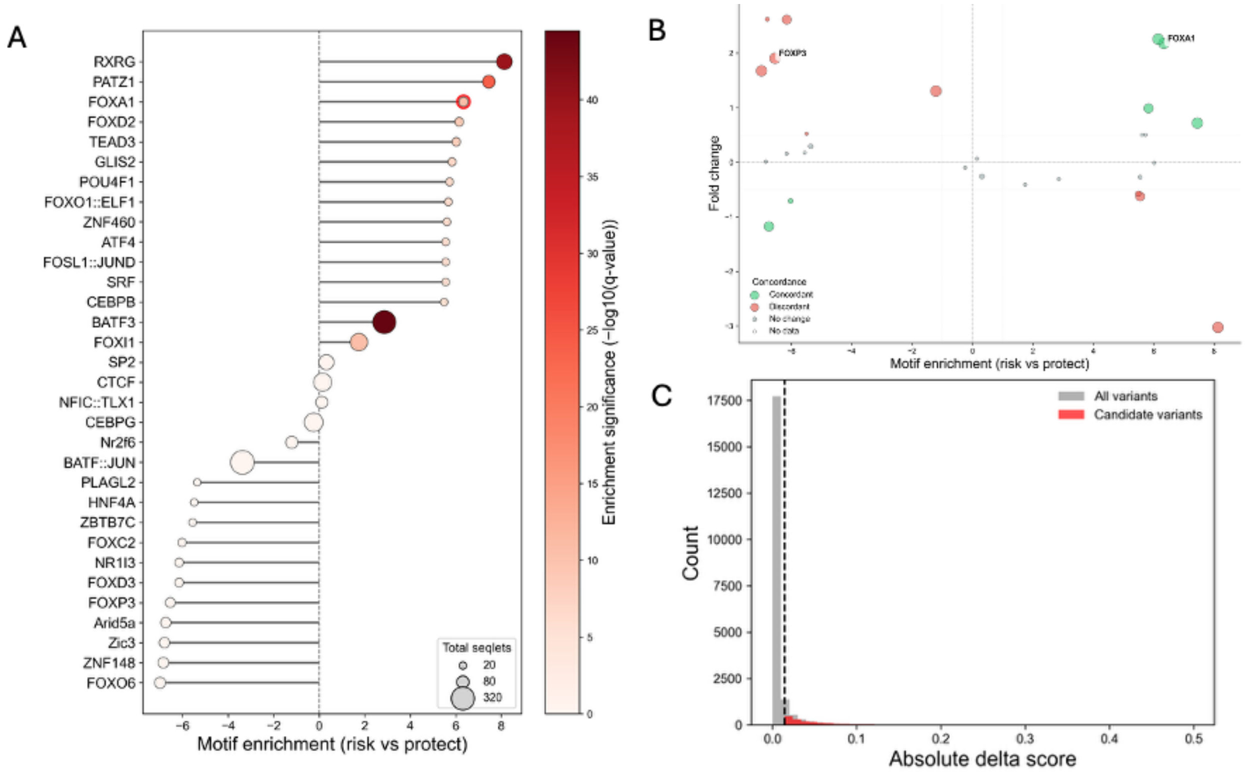


**Figure S2. Allele-dependent motif enrichment and tumor expression concordance in breast cancer.** (A) Transcription factor motifs identified from breast cancer GWAS variants using TREDNet enhancer models and attribution-based TF-MoDISco analysis, ranked by $\log_2$ risk/protective enrichment. Dot size indicates seqlet support; color represents $-\log_{10}$(q-value). (B) Motif enrichment (x-axis; $\log_2$ risk/protective ratio) versus tumor differential expression in TCGA-BRCA (y-axis; $\log_2$ fold change). Concordant TFs (green) show agreement between risk allele enrichment and tumor upregulation. FOXA1 and FOXD2 exhibit strong concordance. Point size reflects $-\log_{10}$(FDR). Dashed lines indicate classification thresholds. (C) Distribution of absolute allele-dependent enhancer prediction differences (|Δscore|) across all variants (gray) and prioritized candidates (red). The dashed vertical line marks the 90th percentile cutoff. Motif enrichment values were computed as $\log_2((n\_risk + 0.5)/(n\_protect + 0.5))$, applying a pseudocount of 0.5 (Haldane-Anscombe correction) to avoid undefined values when the protective allele seqlet count was zero.

**Table S1** Summary of TF motif enrichment and tumor expression. $Log_2$ enrichment values were computed as $log_2((n\_risk + 0.5) / (n\_protect + 0.5))$ using a pseudocount of 0.5 (Haldane-Anscombe correction). Negative $log_{10}$(q-values) are reported for statistical significance.

| TF motif | N-seqlets protect allele | N-seqlets risk allele | $log2^{enrich}$ (risk vs. protect) | $neglog10^q$ | Gene | logFC | FDR |
|---|---|---|---|---|---|---|---|
| BATF3 | 38 | 279 | 2.86 | 44.50 | BATF3 | -0.31 | 0.00 |
| RXRG | 0 | 140 | 8.13 | 39.58 | RXRG | -3.03 | 0.00 |
| PATZ1 | 0 | 87 | 7.45 | 23.91 | PATZ1 | 0.72 | 0.00 |
| FOXI1 | 40 | 135 | 1.74 | 10.61 | FOXI1 | -0.41 | 0.10 |
| FOXA1 | 0 | 40 | 6.34 | 10.50 | FOXA1 | 2.18 | 0.00 |
| FOXD2 | 0 | 35 | 6.15 | 9.16 | FOXD2 | 2.26 | 0.00 |
| TEAD3 | 0 | 32 | 6.02 | 8.37 | TEAD3 | -0.01 | 0.87 |
| GLIS2 | 0 | 28 | 5.83 | 7.29 | GLIS2 | 0.99 | 0.00 |
| POU4F1 | 0 | 26 | 5.73 | 6.77 | POU4F1 | 0.50 | 0.29 |
| FOXO1::ELF1 | 0 | 25 | 5.67 | 6.53 | - | - | - |
| ZNF460 | 0 | 24 | 5.61 | 6.29 | ZNF460 | 0.50 | 0.98 |
| ATF4 | 0 | 23 | 5.55 | 6.11 | ATF4 | -0.27 | 0.00 |
| FOSL1::JUND | 0 | 23 | 5.55 | 6.11 | - | - | - |
| SRF | 0 | 23 | 5.55 | 6.11 | SRF | -0.63 | 0.00 |
| CEBPB | 0 | 22 | 5.49 | 5.86 | CEBPB | -0.58 | 0.00 |
| SP2 | 57 | 71 | 0.31 | 0.26 | SP2 | -0.26 | 0.00 |
| CTCF | 90 | 100 | 0.15 | 0.01 | CTCF | 0.06 | 0.11 |
| PLAGL2 | 20 | 0 | -5.36 | 0.00 | PLAGL2 | 0.29 | 0.00 |
| ZBTB7C | 23 | 0 | -5.55 | 0.00 | ZBTB7C | 0.17 | 0.24 |
| ZNF148 | 57 | 0 | -6.85 | 0.00 | ZNF148 | 0.01 | 0.90 |
| FOXP3 | 46 | 0 | -6.54 | 0.00 | FOXP3 | 1.91 | 0.00 |
| NR1I3 | 35 | 0 | -6.15 | 0.00 | NR1I3 | 0.16 | 0.09 |
| NFIC::TLX1 | 37 | 40 | 0.11 | 0.00 | - | - | - |

| | | | | | | | |
|---|---|---|---|---|---|---|---|
| HNF4A | 22 | 0 | -5.49 | 0.00 | HNF4A | 0.52 | 0.00 |
| FOXO6 | 63 | 0 | -6.99 | 0.00 | FOXO6 | 1.67 | 0.00 |
| FOXD3 | 35 | 0 | -6.15 | 0.00 | FOXD3 | 2.61 | 0.00 |
| FOXC2 | 32 | 0 | -6.02 | 0.00 | FOXC2 | -0.71 | 0.00 |
| CEBPG | 110 | 93 | -0.24 | 0.00 | CEBPG | -0.10 | 0.07 |
| BATF::JUN | 305 | 29 | -3.37 | 0.00 | - | - | - |
| ARID5A | 53 | 0 | -6.74 | 0.00 | ARID5A | -1.18 | 0.00 |
| NR2F6 | 56 | 24 | -1.21 | 0.00 | NR2F6 | 1.30 | 0.00 |
| ZIC3 | 55 | 0 | -6.79 | 0.00 | ZIC3 | 2.62 | 0.00 |

**Table S2** Enrichment of approved drugs among candidate compounds at varying thresholds. P-values were calculated using Fisher's exact test.

| Percentile | N-candidate | N-approve | Enrichment fold | Odds ratio | P value |
|---|---|---|---|---|---|
| 0.75 | 157 | 24 | 1.10 | 1.13 | 0.33 |
| 0.8 | 125 | 21 | 1.21 | 1.26 | 0.20 |
| 0.85 | 94 | 18 | 1.38 | 1.48 | 0.09 |
| 0.9 | 63 | 17 | 1.95 | 2.32 | 0.00 |
| 0.95 | 32 | 6 | 1.35 | 1.44 | 0.28 |

**Table S3** Compound names corresponding to the column order shown from left to right in Figure 4B, C

| Z score based | Spearman based |
|---|---|
| ixazomib | ixazomib |
| pitavastatin | fulvestrant |
| pralatrexate | pitavastatin |
| fulvestrant | pralatrexate |
| irinotecan | irinotecan |
| bortezomib | reserpine |
| carfilzomib | perampanel |
| perampanel | pentobarbital |
| pentobarbital | bortezomib |
| amsacrine | carfilzomib |
| reserpine | amsacrine |
| homosalate | mycophenolate-mofetil |
| bisacodyl | thiothixene |
| mycophenolate-mofetil | homosalate |
| clofarabine | bisacodyl |
| ingenol-mebutate | ingenol-mebutate |
| thiothixene | clofarabine |
| floxuridine | floxuridine |
| BRD-K78385490 | cercosporin |
| BRD-K69894866 | BRD-K78385490 |
| BNTX | BRD-K69894866 |
| cercosporin | BVD-523 |
| BRD-A40431293 | BRD-K60870698 |
| NNC-55-0396 | BRD-K28366633 |
| BRD-K74316684 | NSC-3852 |
| JNJ-26481585 | JNJ-26481585 |
| genz-644282 | BRD-A40431293 |
| camptothecin | NNC-55-0396 |
| BRD-A49848186 | CVF-SUMO-11 |
| NSC-3852 | ST-056792 |
| BRD-K28366633 | BVT-948 |
| AG-592 | EMF-sumo1-39 |
| malonoben | AG-592 |
| BRD-K60870698 | VU-0418934-2 |
| AMG-232 | BRD-A49848186 |
| BVD-523 | BRD-K55722623 |

| | |
|---|---|
| VU-0418934-2 | BRD-K74305673 |
| RG-7388 | BRD-K81795824 |
| BRD-K74305673 | BRD-A68065211 |
| BRD-K81795824 | malonoben |
| BRD-K52321331 | BRD-K74316684 |
| CVF-SUMO-11 | BNTX |
| BRD-A68065211 | BRD-K52321331 |
| ST-056792 | BRD-K95285735 |
| PHA-848125 | AMG-232 |
| EMF-sumo1-39 | camptothecin |
| BRD-A37735495 | BRD-K18724229 |
| BRD-K18724229 | deguelin |
| merck-ketone | PHA-848125 |
| deguelin | BRD-A49680073 |
| BRD-K55722623 | hycanthone |
| BVT-948 | merck-ketone |
| YM-155 | brazilin |
| BRD-A49680073 | genz-644282 |
| hycanthone | RG-7388 |
| BRD-K95285735 | YM-155 |
| R-547 | SB-939 |
| brazilin | R-547 |
| dorsomorphin | dorsomorphin |
| diphenyleneiodonium | BRD-A37735495 |
| BRD-K00313977 | BRD-K00313977 |
| SB-939 | diphenyleneiodonium |
| BRD-K18726304 | BRD-K18726304 |

**Table S4** Number and fraction of candidate compounds with positive pathway anti-correlation scores and Spearman correlation (spr) across pathways.

| pathway | N-positive | Positive (%) | N-positive(spr) | Positive-spr (%) |
|---|---|---|---|---|
| G2M CHECKPOINT | 63 | 100 | 63 | 100 |
| E2F TARGETS | 62 | 98.4 | 62 | 98.4 |
| ESTROGEN RESPONSE LATE | 62 | 98.4 | 61 | 96.8 |
| MTORC1 SIGNALING | 62 | 98.4 | 61 | 96.8 |
| PI3K AKT MTOR SIGNALING | 60 | 95.2 | 58 | 92.1 |
| MYC TARGETS V1 | 59 | 93.6 | 58 | 92.1 |
| UNFOLDED PROTEIN RESPONSE | 57 | 90.5 | 52 | 82.5 |
| ESTROGEN RESPONSE EARLY | 44 | 69.8 | 48 | 76.2 |
